# Comparison of NMR and crystal structures of membrane proteins and computational refinement to improve model quality

**DOI:** 10.1101/127142

**Authors:** Julia Koehler Leman, Andrew R. D’Avino, Yash Bhatnagar, Jeffrey J. Gray

## Abstract

Membrane proteins are challenging to study and restraints for structure determination are typically sparse or of low resolution because the membrane environment that surrounds them leads to a variety of experimental challenges. When membrane protein structures are determined by different techniques in different environments, a natural question is “which structure is most biologically relevant?” Towards answering this question, we compiled a dataset of membrane proteins with known structures determined by both solution NMR and X-ray crystallography. By investigating differences between the structures, we found that RMSDs between crystal and NMR structures are below 5 Å in the membrane region, NMR ensembles have a higher convergence in the membrane region, crystal structures typically have a straighter transmembrane region, have higher stereo-chemical correctness, and are more tightly packed. After quantifying these differences, we used high-resolution refinement of the NMR structures to mitigate them, which paves the way for identifying and improving the structural quality of membrane proteins.

## Introduction

Membrane proteins (MPs) are important drug targets^1^, yet remain extremely difficult to study because of the hydrophobic environment that surrounds them. This situation leads to a number of questions, some of which we attempt to investigate here: How well do membrane mimetics (see Supplement) used for structure determination mimic native membranes? How do MP structures differ from their soluble counterparts? How many of these differences can be attributed to (1) the hydrophobic environment of the membrane; (2) the chosen method for structure determination; (3) the different environments in a crystal vs. solution; (4) the influence of the membrane mimetics; and (5) the different environment in specific lipids or detergents? How can we identify and quantify ‘artifacts’ arising during structure determination? And how can we correct such features?

The membrane bilayer is a heterogeneous environment composed of chemically different lipids^2,3^ that are in constant movement around the protein. An asymmetric distribution between the inner and outer leaflet^4^, lateral diffusion, and formation of ordered micro-domains (i.e. lipid rafts^5^) make the natural bilayer difficult to model experimentally or computationally. Further, to determine MP structures, the protein must be concentrated in a chosen membrane mimetic, that often depends on the method for structure determination, where the exact conditions are found through trial and error^6,7^. Crystallographers often use micelles or lipidic cubic phases^8^, optimizing conditions based on crystal quality and diffracting resolution, typically sacrificing a native-like environment. Similarly, for NMR spectroscopy ‘optimal conditions’ are found by improving spectral quality to obtain the largest number of restraints, which, for MPs, are often sparse and of low resolution. These restraints are then used to compute structural models via MD simulations, which in turn work best with rich restraint datasets that are typically difficult to obtain for MPs^9^. In comparison to crystallography where protein flexibility often leads to missing density and hence to missing atom coordinates, a lack of restraints in NMR does not lead to missing atom coordinates, but rather to less reliable ones. The latter poses a problem when a single NMR model is used as a representative of an entire ensemble, which is unfortunately done too often.

One can therefore anticipate that some of the differences between NMR and crystal structures arise from the method for structure determination. For soluble proteins, Garbuzynskiy and co-workers compiled a set of 78 proteins^10^ and found that the RMSD between NMR and crystal structures are generally larger than the RMSD within the NMR ensemble, which indicates well-converged NMR ensembles. Montelione and co-workers came to similar conclusions on 148 structure pairs^11^, where for 76% of the pairs this difference was larger than a factor of two. On average, the backbone RMSD between crystal and NMR structure was about 1.0 and 1.4 Å over core and all residues, respectively, and was independent of age or quality of the NMR models.

Carugo et al. saw similar RMSDs (in the range of 1.5-2.5 Å) on 109 soluble protein structure pairs^12^. They found that sidechain conformations of hydrophobic amino acids were more similar in crystal and NMR structures than of hydrophilic ones, which is expected because hydrophobic amino acids are well-packed and buried in the protein interior. Further, differences between NMR and crystal structures were on average smaller for β-strands than for helix or loop residues, differences in loops were independent of crystal packing effects, and rotamer variations of buried sidechains were rare.

Here, we identify measures to quantify structural differences in MPs determined by crystallography and NMR. So far, there are no well-established metrics to measure the biological validity of a MP structure, and experimental verification remains difficult as structure determination in native-like membranes or vesicles remains virtually impossible to date. Functional assays can verify biological activity of the protein, yet, these are often carried out in vesicles or cells under very different conditions than those being used for structure determination^13^. Conversely, functional assays in membrane mimetics used for structure determination (micelles, bicelles or nanodiscs) are practically challenging. Determining structures in multiple mimetics with different techniques provides insights, but is extremely labor intensive and costly. Nevertheless, this has been accomplished for some proteins^13^, for instance the influenza virus A M2 protein^14^ and bacteriorhodopsin.

It was suggested that, once non-native features are identified in a MP structure, computational methods, such as restrained molecular dynamics^15,16^, may be used for structural refinement^13^. Lower-cost alternatives are Monte-Carlo simulations, for instance in the Rosetta software suite. For soluble proteins, Mao et al. used restrained refinement and *de novo* structure prediction in Rosetta in conjunction with NMR data to refine NMR models^17^. Their refined models had smaller RMSDs to the crystal structures and were better suited for phasing them. Further, the refined models had on average fewer dihedral angle restraint violations and had improved stereochemistry.

Here we investigate the differences between NMR and crystal structures of MPs on a dataset of 14 structure pairs; to our knowledge, this is the largest comparison of MP structures so far. We use this dataset to establish measures to evaluate the structural quality, identify differences from structure determination, and test an efficient computational refinement approach in Rosetta to reduce them. While questions about the influence of distinct membrane mimetics and lipid specificity remain due to the small number of currently available structure pairs, we provide measures to identify and correct for non-native features in MP structures.

## Results and Discussion

We curated a dataset of MP structure pairs determined by both crystallography and solution state NMR. (Since solid-state NMR structures don’t exist for all of the proteins in the database and only very few cases exist for M2 and DsbB, they were excluded.) We chose the pairs according to resolution, convergence in the NMR ensemble, and sequence similarity (with fewest mutations as possible) and removed all ligands and hetero-atoms (see Methods). The final dataset contained seven α-helical and seven β-barrel MPs with protein sizes from 100 to 363 residues (Table 1 and Figure 1). While this dataset is too small to derive information of statistical significance, it is an excellent starting point to identify differences and trends within those structure pairs that can be substantiated when more structures become available.

**Table 1.**
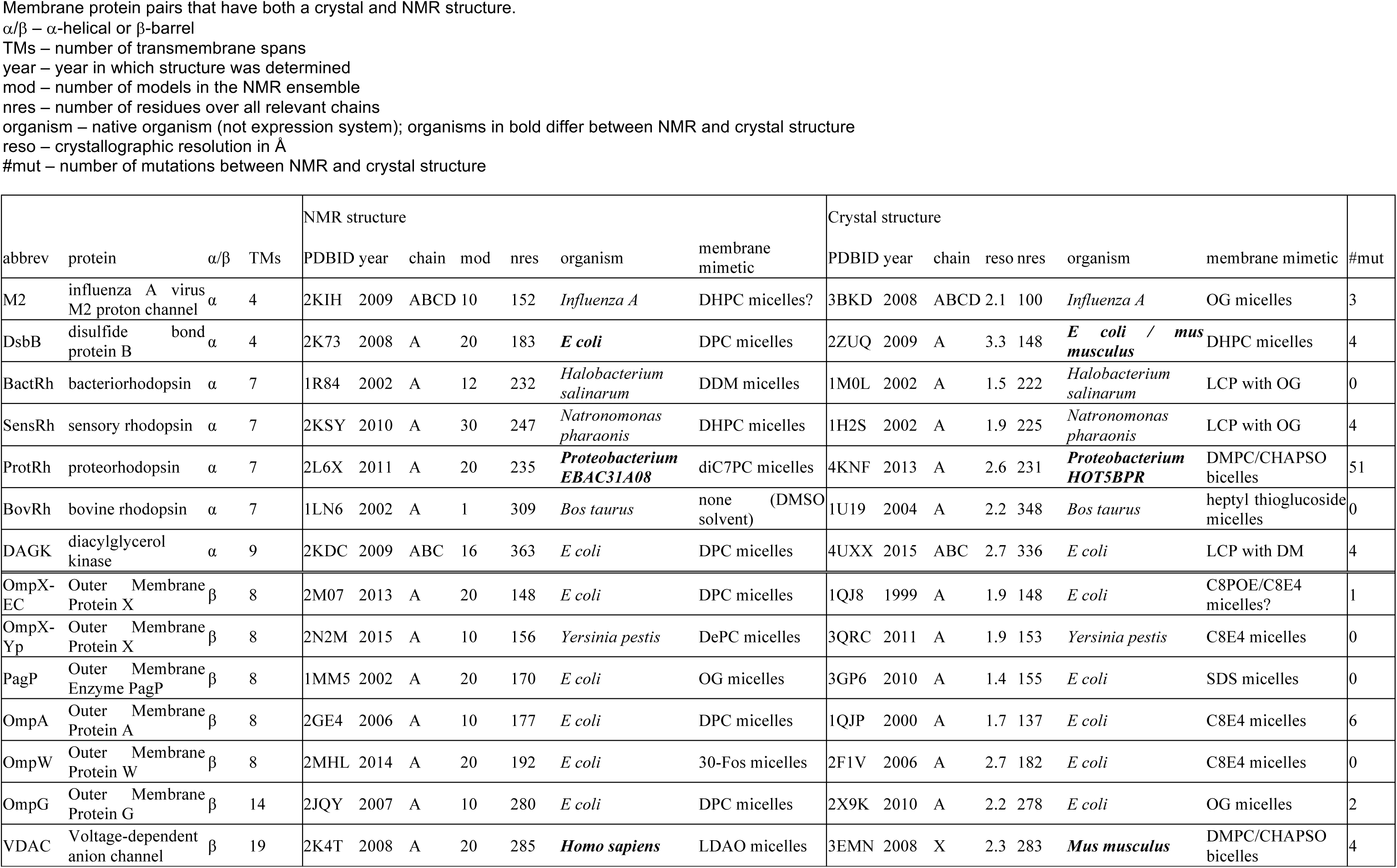
Membrane protein pairs that have both a crystal and NMR structure.

**Figure 1.**
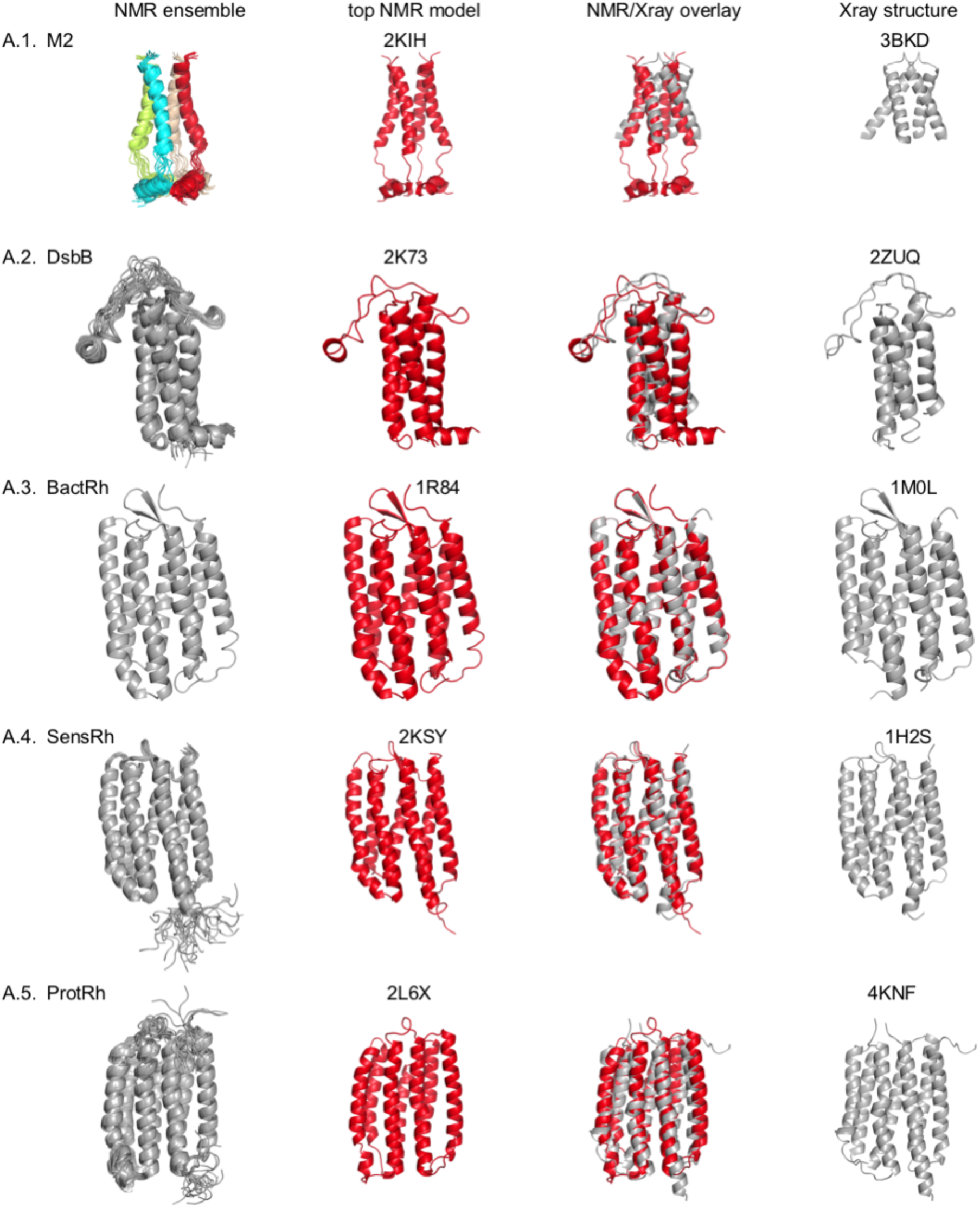
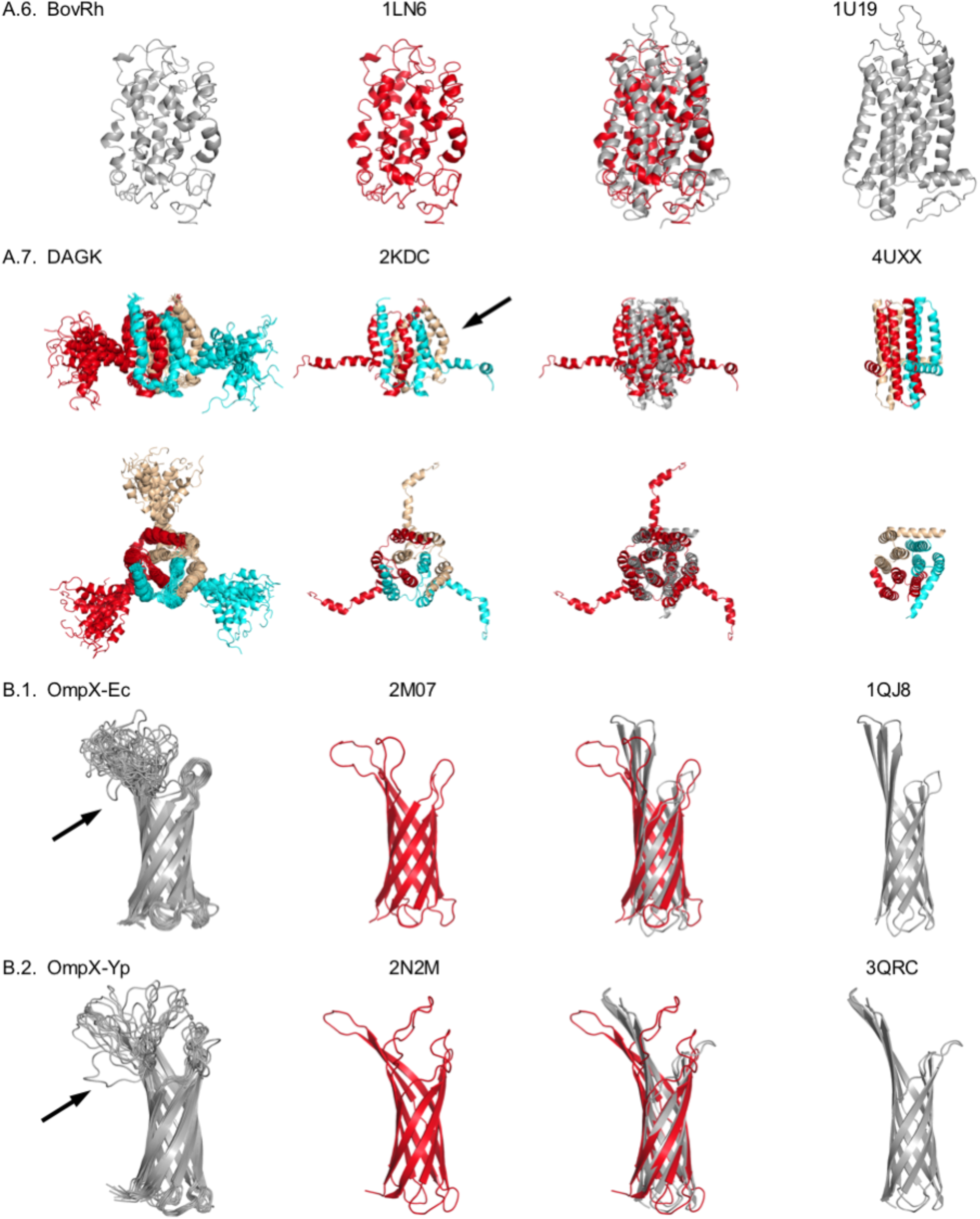
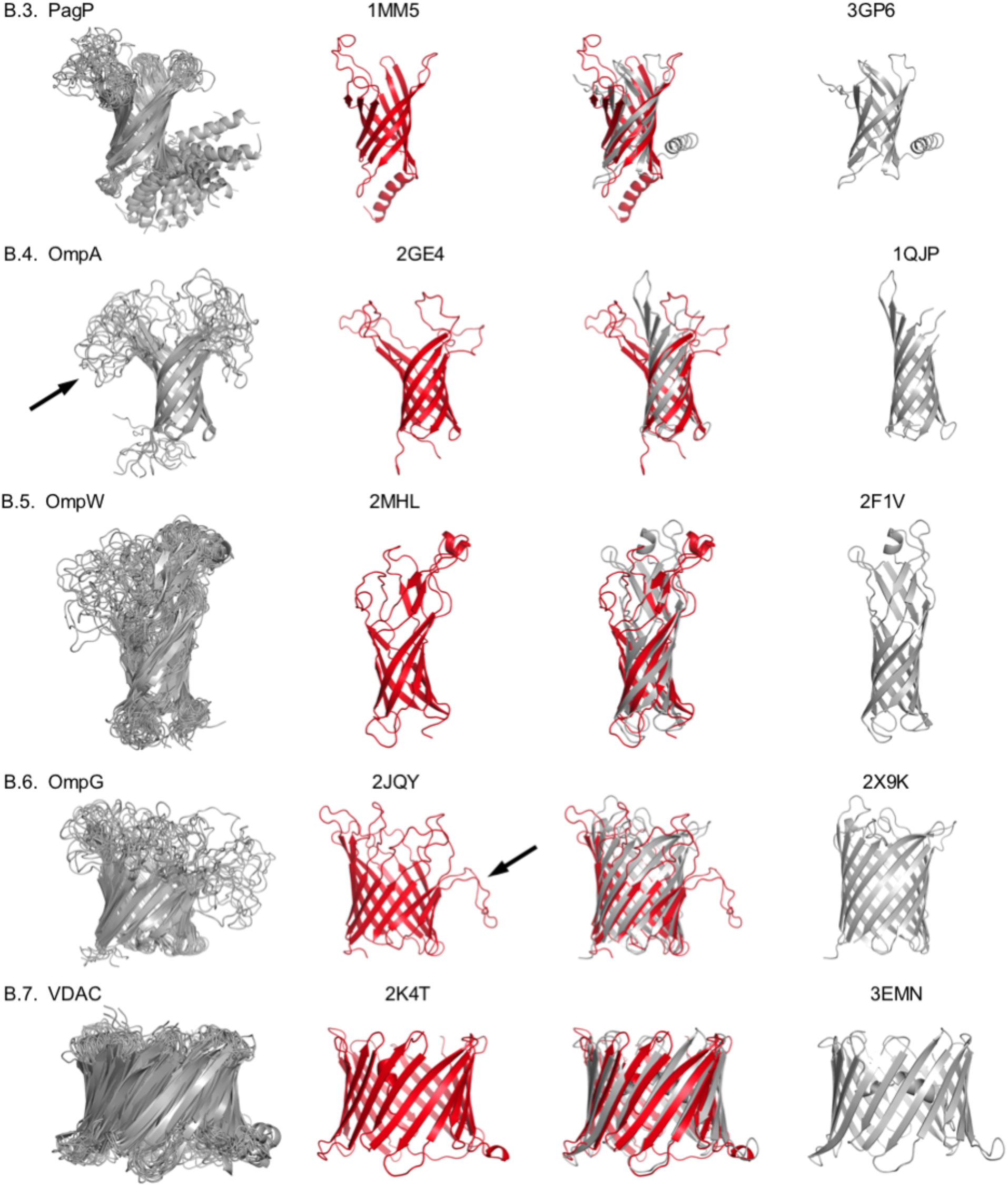
Membrane protein pairs determined by both NMR and Xray crystallography. Proteins are ordered by size, in α-helical and β-barrel categories. The columns from left to right show the NMR ensemble, the top NMR model, the overlay between NMR and crystal structure (red – NMR; gray - Xray), and the crystal structure. Black arrows in DAGK, OmpX, OmpA, and OmpG identify bends in the structures, mostly through loop regions that wrap around the micelle in which the structure was determined.

The biggest differences between NMR and crystal structures occur for (1) DAGK, which has a domain swap in the NMR structure that is not present in the crystal structure (more on this below). (2) The NMR structure of bovine rhodopsin (BovRh) is one of the earliest structures computed from overlapping fragments determined in DMSO – this structure was added for completeness purposes. (3) M2, which has different functional states with a different opening angle. All crystal structures have a large pore opening and all NMR structures have a small pore opening, which impacts superposition. Both structures are also from different constructs (see Figure 1).

### The TM regions of NMR ensembles converge within 5 Å RMSD

For NMR structures, convergence is typically used as a measure of structural quality. To estimate convergence of an NMR ensemble, we computed backbone RMSDs between the top NMR model to all other models in the ensemble. We treat the top model as the baseline as it has been identified as the lowest-energy conformer during structure calculations via MD simulations. Figure 2 shows that the backbone RMSDs over all residues extends as high as ~14 Å due to regions in the protein that are not well-restrained. We see such less restrained regions for the amphipathic helices in both DAGK and PagP (Figures 1 A.7 and B.3) and for the long, unstructured C-termini of DsbB and SensRh (Figures 1 A.2 and A.4); unstructured termini were excluded for these calculations (see Methods). Further, long loops in OmpA, OmpW, and OmpG give rise to RMSDs up to ~10 Å (Figures 1 B.4-6). When considering only TM regions, the backbone RMSDs are below ~5 Å for all proteins with the largest ranges for VDAC, OmpW, and DAGK. The coordinates of BactRh have an extraordinarily high convergence, with all 12 NMR models being within 0.1 Å RMSD!

**Figure 2.**
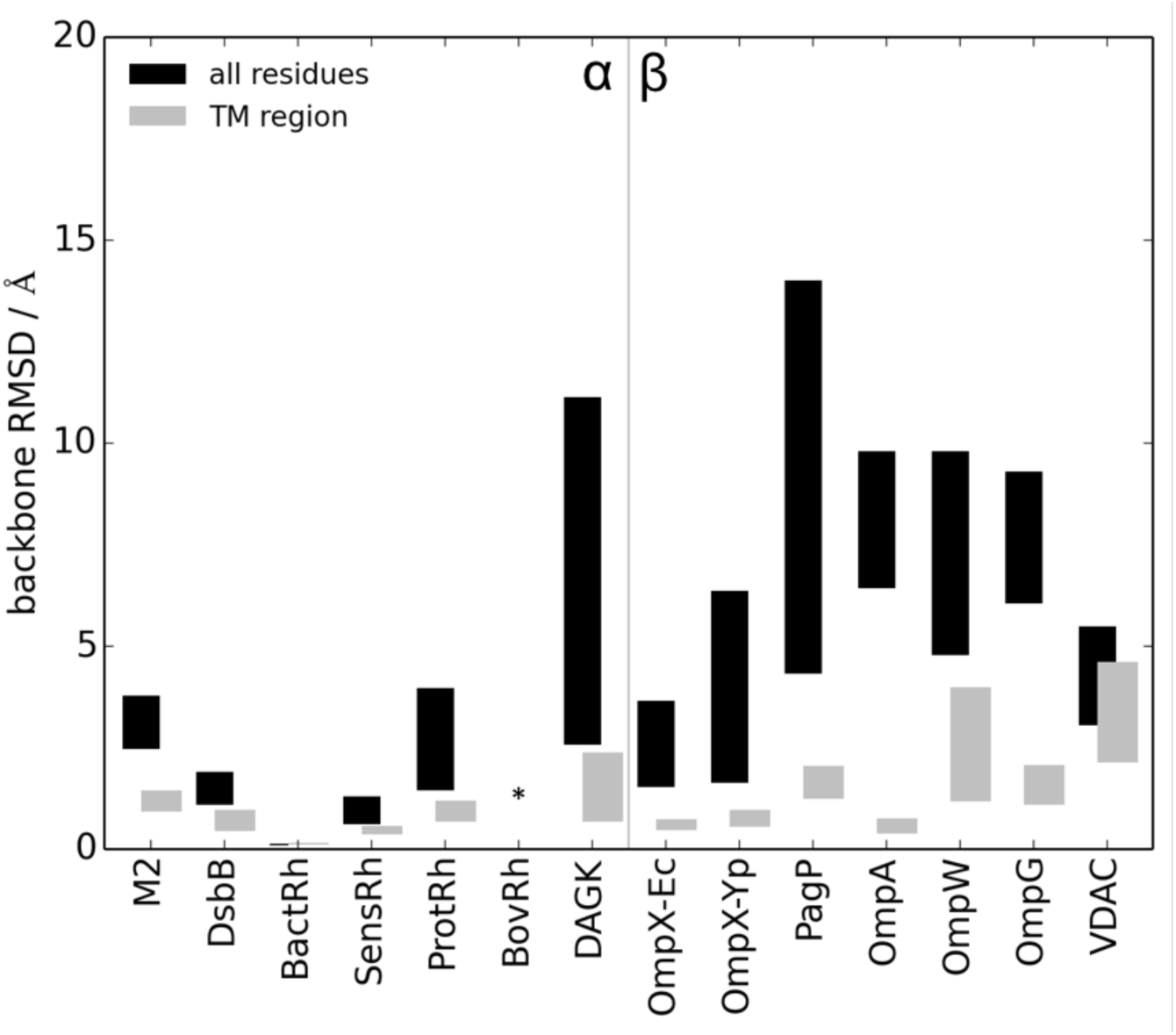
Backbone RMSDs within the NMR ensemble are an indicator of convergence. Ranges of backbone RMSDs between the top NMR model and all other models in the ensemble. RMSDs are computed over all residues (black), and over the TM region (gray) for the α-helical and β-barrel proteins ordered with increasing size and separated by a vertical line. Long, ‘flexible’ C-termini of DsbB (19 residues) and sensory rhodopsin (21 residues) were removed for these calculations. The average backbone RmSd is 4.0 Å over all residues and 1.0 Å for the TM region. The PDB file for bovine rhodopsin only contains a single model and therefore an RMSD cannot be calculated (* asterisk in Figure).

### Convergence in the NMR ensemble is required but insufficient to satisfy model quality measures

So how does convergence of the NMR ensemble relate to ‘structural quality’? To answer this question, we need to distinguish between (1) the convergence *within* the NMR ensemble and (2) the differences *between* NMR and crystal structures. First, convergence is a measure that describes the NMR ensemble, i.e. the relation of different models with respect to each other (completely independent of the crystal structure). Convergence simply means that there is a sufficient number of restraints to create models with little variation between them. Hence there is higher convergence in the TM region due to the larger number of measured restraints. Further, there are no enforced standards on how many NMR models are reported in the PDB and how many models are computed. For instance, the top 10 structures from 1000 computed trajectories might have a better convergence, yet expose subtleties of the energy function used in the calculation, while the top 10 structures from 50 generated models might show more structural diversity.

Secondly, the NMR and crystal structures are almost always determined under different environmental conditions, which can give rise to small structural variations. That means that a well-converged NMR ensemble can still differ from the crystal structure, as we can see in the splaying of OmpA (Figure 1 B.4). Also, the crystal structure of *E. coli* OmpX shows a more elliptic barrel, while it is more circular in the well-converged NMR ensemble (data not shown). These differences might also be explained by the type of restraints used for NMR model calculations. For β-barrels, NMR restraints from backbone Nuclear Overhauser Enhancements (NOE’s, i.e. distance restraints up to 6 Å between atoms more than 4 residues apart) used for structure calculations are long-range restraints in terms of sequence separation, while they are local restraints considering the measured distance. However, (in contrast to crystallographic electron density maps) their long-range character is insufficient to reach across the barrel to distinguish between a circular or elliptic barrel. Further, differences between NMR and crystal structures may also originate in the fact that MD simulation methods used for structure calculations are not designed for sparse datasets, and tools that are better suited are not widely used for structure calculations. For example, the RASREC protocol in Rosetta has been shown to produce higher-quality solution structures with sparse data^9,18^.

Interestingly, we observe a correlation (R = 0.84) between the maximum backbone RMSD within the NMR ensembles and the backbone RMSDs between the crystal and NMR structures (Supplemental Figure 4), when DAGK is removed because of its domain swap, the correlation is as high as 0.90. This correlation likely arises because atoms, that have large deviations from the average model within the NMR ensemble, also deviate considerably from the crystal structure (i.e. these would be weakly restrained regions).

### The type of NMR restraints influences structural quality

Similarly, as the structure determination method can influence the structural representation, the number and types of NMR restraints can influence the quality of the NMR structure. Additionally, which becomes especially important for large proteins like MPs, the type of the protein (α-helical bundle or β-barrel) influences the type of measurable restraints as well as the type of restraints required for protein fold determination or structure calculation. For β-barrels, the resonances in the NMR spectra have a larger spectral dispersion, leading to less signal overlap. This facilitates and therefore increases the number of possible resonance assignments and the acquisition of restraints. Further, protein fold determination for β-barrels is possible through backbone hydrogen bonds between neighboring strands, which requires ‘as little as’ backbone assignments and the measurement of long-range NOEs between neighboring strands. For α-helical proteins however, the spectral dispersion is reduced, which results in larger resonance overlap, complicating and therefore decreasing the number of resonance assignments and hence the number of measured restraints. It is much more difficult to obtain sidechain assignments and restraints involving sidechain atoms between neighboring α-helices, which are especially important for protein fold determination of α-helical proteins. Including long-range restraints from sidechain atoms in structure calculations can drastically improve structural quality, as has been shown for SensRh, for which over 1500 long-range sidechain restraints could be measured via ILV methyl labeling (Supplemental Table 1). If such NOEs are unavailable for helical bundles, PREs or RDCs are commonly acquired to compensate for the lack of long-range restraints. In addition to the difference in measurable restraints and structure calculation, no standard format exists on how structural quality measures are reported, even if NOEs are reported. For instance, when gathering information about the published structures for Supplemental Table 1, NOEs for α-helical proteins were reported separately for intra-residue, sequential, medium-range and long-range NOEs. For β-barrels, even though most of the NOEs are long-range (see VDAC in Supplemental Table 1), no such distinction was made for most of the proteins.

For the α-helical structures in our database, SensRh has a well-defined ensemble with the largest number of long-range NOEs per residue. For the β-barrels, the structure can be determined by a comparably small number of long-range NOEs alone: For OmpX, the *yersinia pestis* construct has about three times the number of long-range NOEs as the *E coli* construct, yet the backbone RMSDs in the TM region are similarly small. OmpA is the only barrel for which RDCs were measured and the TM domain has the best convergence among β-barrels, as indicated by the smallest backbone RMSDs within the ensemble.

### TM regions between crystal and NMR structures agree within 5 A RMSD

To identify differences between NMR and crystal structures, we computed the backbone RMSDs between them for trimmed models that had the same number of residues (see Methods). The backbone RMSDs for the full-length structures can vary considerably (Figure 3A). DAGK contains a domain swap of the helices between the NMR and crystal structures^19,20^ (see below for discussion on this); the RMSD over the TM regions without the domain swap is 6.9 Å. Large differences are also seen for PagP (Figure 1 B.3), which contains an amphipathic N-terminal helix (which is either dynamic, poorly restrained, or both – see below), and OmpG (Figure 1 B.6), which has a long loop that protrudes straight in the crystal structure while wrapping around the micelle in the NMR model. Further, the NMR model of BovRh is distorted since it was computed from structures of overlapping fragments in DMSO solvent. For M2, the structural alignment is inferior due to opening angle differences between NMR and crystal structures that comes from different functional states. Interestingly, for structures containing the TM domain, all crystal structures have a larger opening angle and all NMR (even solid-state NMR) structures have a smaller pore.

**Figure 3.**
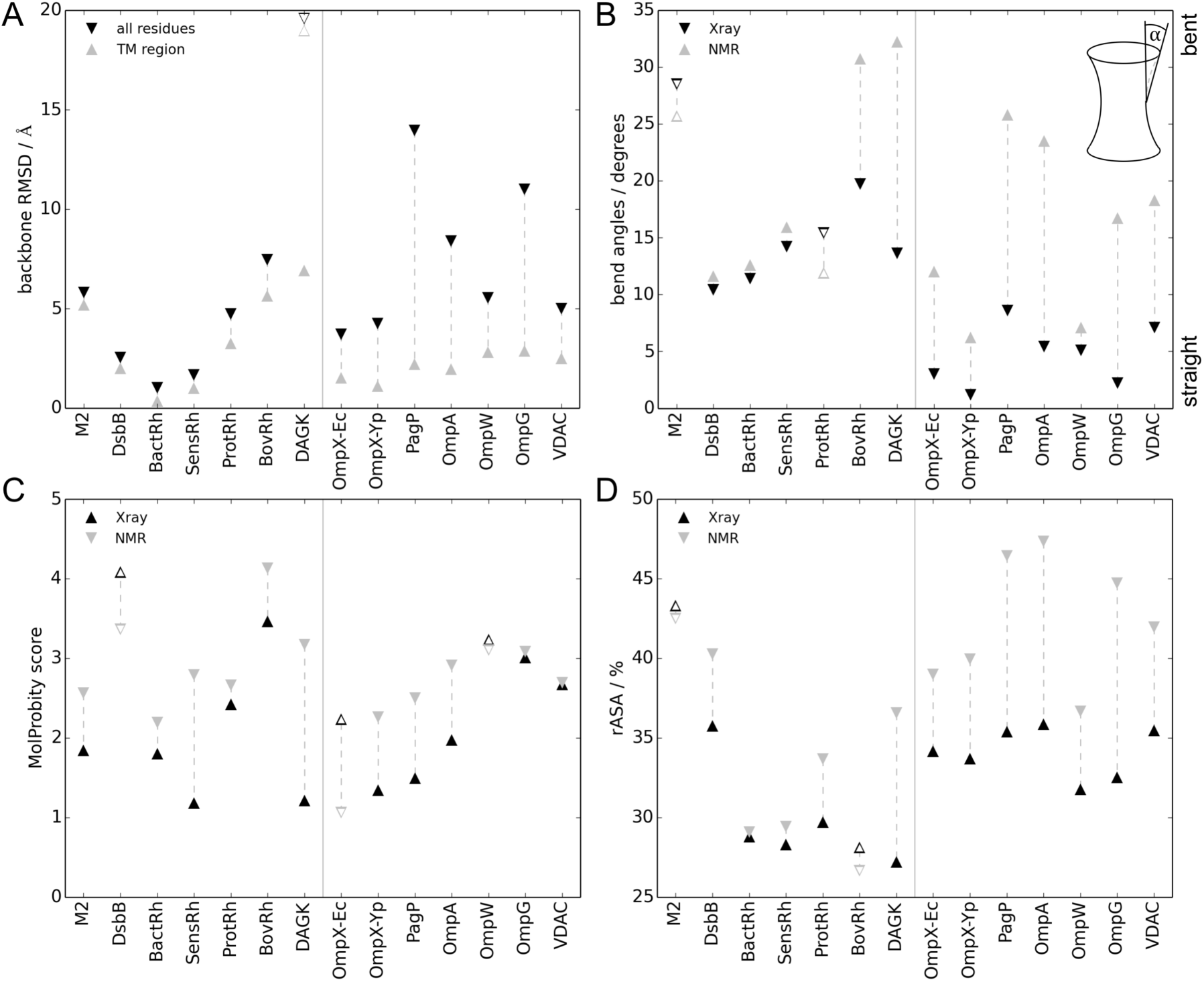
Differences between NMR and crystal structures according to different quality measures. (A) Backbone RMSDs between NMR and crystal structures. The backbone RMSDs between the top NMR model and the crystal structure are computed over all residues (black) and over the TM region (gray). DAGK is an outlier due to a domain swap between the crystal and NMR structure (open symbols); the backbone RMSD for the TM region without the domain swap is 6.9 Å (closed symbols). Except for DsbB, the helical proteins contain short loops which are rather rigid and hence the difference between RMSDs of the full-length protein and the TM region is small. The β-barrels have long, ‘flexible’ loops that give rise to a larger RMSD in the full-length structure compared to the TM region. For β-barrels the RMSDs in the TM region vary little among proteins, likely because there is little flexibility in the barrel region itself that is stabilized by an extensive hydrogen bonding network involving backbone atoms. (B) Crystal structures typically have a straighter TM region than NMR structures. We use bend angles as a straightness measure for the protein TM regions – the definition of the bend angle is shown in the lower right corner of the plot. Bend angles of the crystal structures are shown in black, for the NMR structures in gray. The hollow triangles deviate from the general trend that crystal structures typically have smaller bend angles, i.e. a straighter TM region, than the NMR structures. (C) Crystal structures typically have better stereochemistry. We use MolProbity scores as a measure of stereochemistry for the crystal structures (black) and the NMR structures (gray) - lower scores indicate better stereochemistry. The overall MolProbity score combines all individual scores (see Supplemental Table 2) and is a measure of structural quality. The score itself is normalized such that it provides the resolution of the crystal structure at which those values would be expected^24^. Hence, if the MolProbity score is lower than the crystallographic resolution, the quality of the structure is better than the average structure at this resolution. Hollow triangles deviate from the general trend that crystal structures have lower (i.e. better) scores than NMR structures. (D) Crystal structures are typically more tightly packed than NMR structures. We used Naccess to compute the per-residue accessible surface area (ASA) relative to an extended chain, as a measure of packing density. 100% means completely exposed and 0% means completely buried. The per-residue rASA were averaged over the entire protein. Crystal structures (black) and NMR structures (gray) with hollow triangles deviate from the general trend that crystal structures are typically more tightly packed with a smaller average rASA than NMR structures.

Figure 3A further shows that while RMSDs in the TM region vary across α-helical MPs (1.0 – 5.6 Å without DAGK), they are relatively uniform for β-barrels (1.1 – 2.9 Å). The small range for β-barrels suggests little flexibility to satisfy a more rigid backbone hydrogen bond network, which is required to maintain protein stability in the membrane. In contrast, α-helical proteins may permit more flexibility in their TM region due to the irregular hydrogen bond network among sidechain atoms, in which bonds can be broken and reformed between different residues without compromising overall protein stability in the membrane.

For full-length constructs, backbone RMSDs between crystal and NMR structures are influenced by loop regions that are either dynamic, poorly restrained, or both. As seen from Figure 3A, backbone RMSDs between the full-length proteins and the TM regions differ substantially for β-barrel proteins (2.2 – 11.8 Å) due to their long loops. This is not the case for α-helical proteins (0.5 – 1.8 Å difference) in which long loops are absent, except for DsbB whose long loop between TM3 and TM4 has a similar conformation in both the NMR and the crystal structure.

### And what about dynamics?

One of the challenges in comparing NMR and crystal structures is the consideration of dynamics. While crystal structures are static snapshots, the B-factor can indicate thermal motion within an observable range. However, when certain regions are too dynamic, they don’t give rise to well-defined electron density and therefore often have missing atom coordinates in the crystal structure. NMR spectroscopy can be used to measure dynamics, yet is often interpreted incorrectly: ‘flexible’ regions in an NMR ensemble are generally considered dynamic, which might or might not be true. In reality, ‘flexible’ regions in an NMR ensemble only point to fewer available restraints for these regions, while dynamics can only be measured by relaxation experiments such as T1, T2 or heteronuclear NOEs, which are not captured in the structural ensemble. Due to these inherent differences in methods and interpretations, it remains difficult to address whether ‘flexible’ regions are poorly restrained, truly dynamic, or both, without investigating each region individually.

### Crystal structures more often have smaller bend angles indicating a straight, cylinder-like TM region

While most crystal structures have a straight TM region that resembles a cylinder, some of the NMR structures have TM regions that resemble stretched or compressed cylinders (concave like a one-sheet hyperboloid vs. convex like a wine barrel). Examples are DAGK, OmpA, and OmpG (Figures 1 A.7, B.4 and B.6) – for some of these examples, one can imagine that the loop regions wrap around the surface of the micelle in which the structure was determined. Because such deviations are not present in the crystal structure, we created a bend angle measure to quantify them (see Methods and Figure 3B): a smaller bend angle represents a straighter TM region. Note that the bend angle here defines the general straightness of the entire TM region (see inset in Figure 3B), but is independent of kinks or bends in individual helices or strands.

For many of the proteins in our dataset, the bend angles are 10-20° smaller for the crystal structures, indicating that the TM regions for those structures are straighter. For M2, both NMR and crystal structures splay (the crystal structure slightly more than the NMR structure) which seems an inherent characteristic of the structure. For ProtRh the structural differences are small: while both original PDBs contain the retinal ligand, the crystal structure is more tightly packed, while the NMR structure has a larger ligand cavity.

The largest variations in bend angles (> 10°) can be seen for DAGK, BovRh, PagP, OmpA, OmpG, and VDAC. BovRh is an outlier due to its poor structural quality. DAGK has marked differences in bend angles (Figure 3B), which might be explained in different ways. Proteins crystallize more readily in the absence of dynamics and when they are in a low-energy state, which could be achieved in a dense packing arrangement in the crystal. For DAGK, the tight packing arrangement might lead to a straightening of the helices (see Supplemental Figure 3). Further, the absence of orientational NMR restraints (such as RDCs) that report on helix bending might lead to distortions in the structure. For DAGK, the Q-value^21^ between experimental and back-calculated RDCs is much smaller in the NMR structure (Q=0.03) than for the crystal structure (Q=0.55)., which points to the micellar environment as a major source of differences.

DAGK also has a domain swap in the NMR structure, which is not present in the crystal structure. The domain swap could be explained by slight differences in refolding protocols^22,23^ and/or the ambiguity of NMR restraints used for structure calculation of the homo-oligomeric protein. In fact, Bruce Donald’s group showed that there are nine conformational solutions for DAGK that satisfy the NMR restraints and both the NMR and the crystal structure are valid solutions^20^.

While it is interesting that many (especially β-barrel) structures have larger bend angles in the NMR structure, the origin of these differences remains somewhat elusive (for a summary, see Table 2). Our initial thought was that the smaller lateral pressure in micelles could lead to a less stable environment and therefore to bending. However, many of the crystal structures considered here are also determined in micelles. Another possibility is that loop regions reach out to interact with the curved micelle surface in the NMR structure while they are thermally restricted in the crystal, where they interact with residues of other subunits. Further, crystal structures are generally straighter, irrespective of the membrane mimetic. It is interesting to note that all three structures determined in LCP (BactRh, DAGK, SensRh) and both structures determined in bicelles (ProtRh and VDAC) show lamellar (i.e. stacked) bilayers, whereas all except one structure in micelles (m2) show sponge-like surfaces (Supplemental Figure 3). DsbB has large contacts between the antibody fragments that are inserted for crystallization purposes. Finally, we want to emphasize that while we quantify bend angles here, we don’t want to imply native likeness of one or the other since the membrane mimetics used for structure determination by neither method are biochemically similar to native membranes that occur in the cell.

**Table 2.** Summary of possible influence of the different effects on the MP structure. Effects that are likely larger have more asterisks.

|  | *bend angle:*<br>*large distortions* | *stereochemistry:*<br>*local geometry* | *rASA:*<br>*packing* |
| --- | --- | --- | --- |
| *method:*<br>*Xray vs. NMR*<br>*(direct vs.*<br>*indirect*<br>*detection)* | * bend angles could stem from the fact that loop regions in NMR are less restrained (and we don't know where they really are) | ** NMR restraints during refinement can work against rotamer and Ramachandran dihedrals, so those are worse for NMR, but bond lengths and angles are better in NMR structure because they are refined through the energy function in MD package | less long-range information (across β-barrels) in NMR; error bars in restraints might lead to lower packing density |
| *environment:*<br>*crystal vs.*<br>*solution* | ** crystal packing likely restricts loop conformations | *** crystal environment could lock down rotamers and local geometry | *** protein crystallizes in a low-energy conformation with higher packing density |
| *environment:*<br>*hydrophobic*<br>*membrane*<br>*bilayer* | unclear:<br>solution is isotropic, whereas the membrane is anisotropic | unclear:<br>membrane bilayer could affect observed rotamers | unclear:<br>helix-helix interactions are closer in MPs but entire MPs are less dense than soluble proteins |
| *environment:*<br>*membrane*<br>*mimetic:*<br>*micelle vs.*<br>*bicelle vs. LCP*<br>*etc* | *** loop regions can wrap around soluble micelle, this is energetically unfavorable in a bilayer; bending in DAGK comes from soluble micelle: RDC restraints in DAGK match the NMR structure, but not the Xray structure (not a computational problem) | unclear | ** micelles likely have a smaller lateral pressure than bilayers |
| *environment:*<br>*specific lipids*<br>*/detergents* | unclear:<br>chain length could influence bending to minimize hydrophobic mismatch | unclear | unclear |

### Crystal structures have generally better stereochemistry

To obtain more detailed measures of the structural quality, we used the MolProbity structural validation software^24^. The overall MolProbity score combines measures about steric clashes, Ramachandran outliers and rotamer outliers into a single score; the lower the MolProbity score, the higher the structural quality. Figure 3C shows that the MolProbity scores are typically worse for the NMR structures, except in cases for DsbB, *E. coli* OmpX, and OmpW, with no correlation between the year the structure was determined and its MolProbity score. Supplemental Table 2 contains the individual scores and shows that crystal structures usually have more favored rotamers and Ramachandran dihedrals, while having fewer rotamer and Ramachandran outliers. At the same time, deviations in Cβ positions, bond lengths, bond angles, and proline geometry are often smaller for NMR structures. This seems to indicate that MD simulation software with which NMR structures are computed, is well able to produce good local geometry in bond lengths and angles etc. while it produces less optimal backbone dihedrals and rotamers than in crystal structures. This can occur when the NMR restraints are not turned off (or down-weighted) during the final refinement stage, where they compete with optimal rotamers and Ramachandran angles. Finally, both BovRh and DsbB have bad clash scores for both NMR and crystal structures.

The overall MolProbity clash score (Supplemental Table 2) is similar between NMR and crystal structures. However, the per-residue Rosetta full-atom repulsive score as an indicator of clashes, shows a distinct pattern (Figure 4 and Supplemental Figure 5); While NMR structures typically have more clashes of low severity, crystal structures have few severe clashes. For NMR structures, this may be because a typical criterion for structural quality is satisfaction of restraints from which the model is computed. Considering the large number of required restraints and their typically low quality for MPs, it is difficult to satisfy all restraints at the same time, especially given errors in the data and/or the interpretation of the data, for instance due to incorrect resonance assignments of overlapping peaks. In contrast, in crystallography, the protein structure is traced along the electron density and structure refinement may be carried out with a high degree of automation. It is conceivable that most residues fit the density well but that a few residues are difficult to place due to the low resolution of the electron density and that such errors may easily be overlooked during refinement.

**Figure 4.**
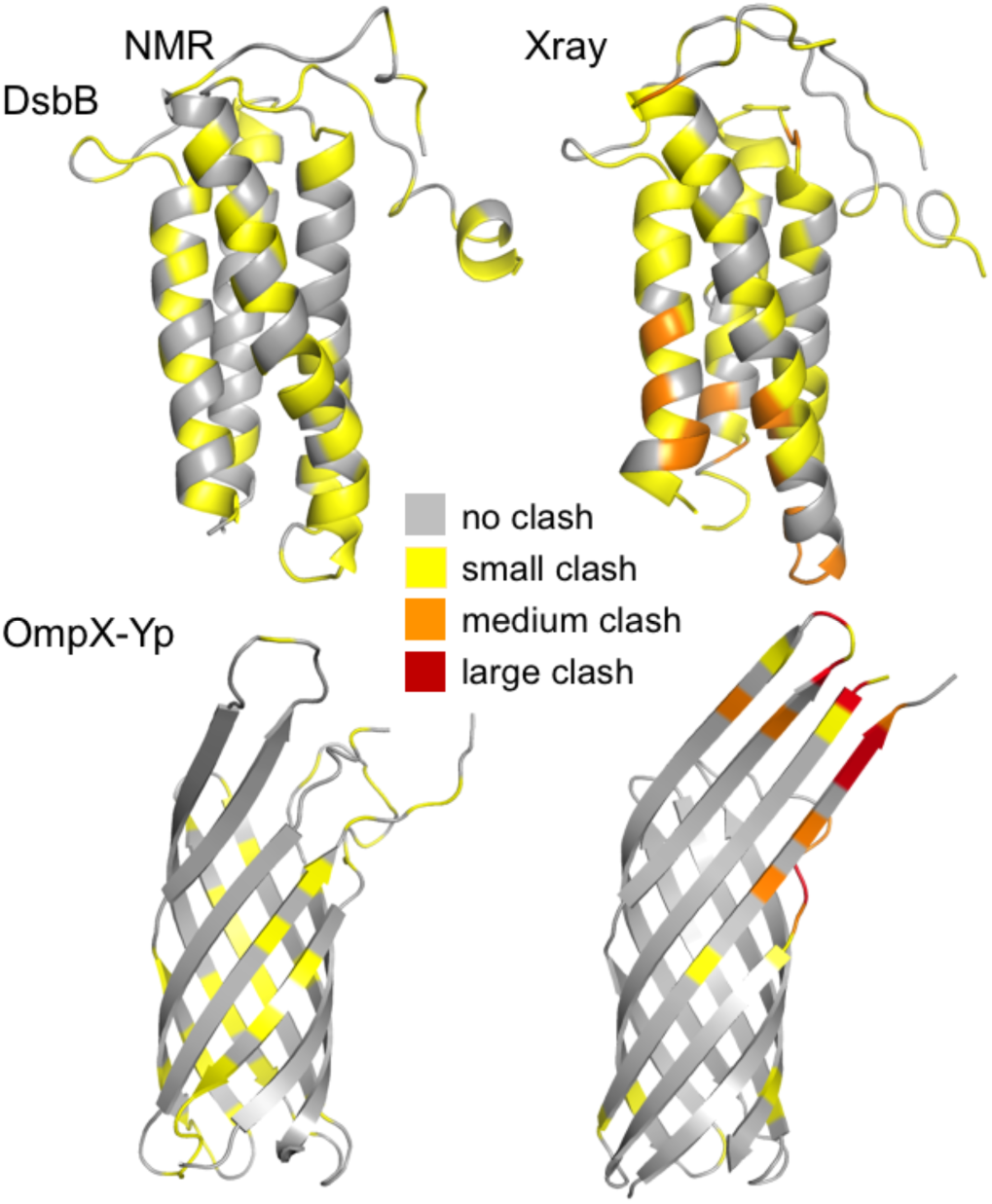
Both NMR and crystal structures contain clashes. Per-residue Rosetta full-atom repulsive scores as an indicator of clashes are mapped onto the protein structures of two examples; the entire dataset is shown in Supplemental Figure 5. The upper panel shows DsbB, the lower panel *Yersinia pestis* OmpX, with the NMR structure on the left and the crystal structure on the right. Residues in gray are clash-free, with increasing clash severity from yellow to orange to red (full-atom repulsive scores <1 REU in gray, 1-10 REU in yellow, 10-100 REU in orange, and >100 REU in red). The results show that NMR structures typically have more clashes, however they are less severe. Crystal structures have fewer clashes, but they are of higher severity.

### Crystal structures generally pack more tightly than NMR structures

We used different measures to probe packing differences between NMR and crystal structures. Figure 3D shows the relative per-residue accessible surface areas (rASA^25^) that are averaged over the number of residues. Except for M2 and BovRh which both have very similar values, the rASA is smaller for crystal structures, indicating that they are more tightly packed. The differences between NMR and crystal structure are usually larger than ~5% rASA.

Tighter packing was also confirmed through calculations of total ASA, protein volume, solvent volume in the protein interior, surface crevices with the 3V software^26^ and packing scores from RosettaHoles^27^ (Supplemental Figure 6). All data reinforce that the NMR structures are less tightly packed, leading to a larger surface area and more rugged surfaces (Supplemental Figure 8) that are more accessible to solvent within the protein and on its surface. We also discuss Glycine accessibility in Supplemental Figure 7.

Tighter packing for crystal structures is consistent with the notion that proteins crystallize more readily in a low-energy state when motion is minimal. Thermal expansion could be considered as the root of packing differences as for all (but one? VDAC?) crystal structures data was collected at 100K while for the NMR structures, spectra were acquired between 283 and 323K. However, this is inconsistent with the fact that for (at least) four MP structures crystallized at different temperatures, the RMSDs between the high- and low-temperature structures were well below 1Å:

○ δ-opioid receptor with PDBIDs 4RWD/4RWA/4N6H at 294/100/100K have an RMSD of 0.8Å
○ adenosine receptor: 5K2C/4EIY at 292/100K have an RMSD of 0.4Å
○ photosystem II: 5WS6/3WU2 at 293/100K have an RMSD of 0.3Å
○ photosynthetic reaction center: 5M7K/2WJN at 293/100K have an RMSD of 0.4Å.

Since thermal expansion is less likely as the source of packing differences, we argue that the different environment between a crystal and solution is a major source of packing differences.

### Packing density could be influenced by the membrane environment

Packing density can further be influenced by the hydrophobic environment of the membrane bilayer. Several groups have examined packing differences between helical proteins in solution and the membrane. When considering structurally similar helix-helix interfaces, MPs pack more tightly because of their large number of small amino acids such as Gly, Ala, and Ser^28,29^. Soluble helix-helix interfaces are often lined by larger hydrophobic residues that cannot make close contacts^30^. In contrast, for whole proteins it was found that MPs pack more loosely than soluble proteins and have more functionally important sites in the cavities^31–33^. These seemingly contrary conclusions may be explained by (1) differing datasets, i.e. structurally similar helix-helix interfaces vs. entire proteins. (2) The hydrophobic effect for soluble proteins leads to tight packing, while the smaller difference in hydrophobicity between the membrane and the hydrophobic interior of the MP (except for pores) may lead to weaker packing. (3) Further, MPs are often transporters and channels with cavities and pores that are functionally relevant and MPs require some structural flexibility to carry out their functions^34^. To investigate the energetics of packing for soluble and MPs, Bowie and coworkers examined the decrease of packing stability with increasing cavity volume between T4-lysozyme and bacteriorhodopsin and found little difference between the two proteins, suggesting similar energetics^35^. However, generalization of this statement requires investigation on a larger database.

### To decipher the influence of membrane mimetics, a larger dataset is needed

Structural differences might also be explained by differences in membrane mimetics (micelles, bicelles, nanodiscs, lipidic cubic phases (LCP)) and lipids/detergents used. While some structures exhibiting marked differences were determined in micelles by both NMR and crystallography (OmpA), other structures exhibiting large differences were determined in different mimetics. For example, the structure of DAGK was determined in detergent micelles by NMR and in lipidic cubic phases by crystallography. Since micelles have increased water penetration in the TM region due to decreased lateral pressure, the structural differences in DAGK likely stem from different membrane mimetic environments^19^. As shown above, back-calculated RDCs from the NMR structure match the NMR structure much better than the crystal structure, which points to the micelle environment as the source of differences. Further, it was shown that predicted resonances from the crystal structure matched data from oriented-sample solid-state NMR much better than for the solution NMR structure^19^, indicating that that liquid crystalline bilayers used in ssNMR are likely similar to the LCP environment in the crystal. For the M2 channel from the influenza A virus a variety of structures is available that were determined in different environments with various techniques (Xray crystallography, ssNMR, solution NMR) for different functional states^36,37^. It is therefore difficult to find a clear correlation between environment (pH, membrane mimetic, specific lipids or detergents). A detailed, statistical analysis of membrane mimetics and an investigation of their influence on structural differences is undoubtedly useful, however difficult to carry out on a large scale due to either lack of data or ambiguity in reporting.

### Identifying non-native features in membrane protein structures

Related to the question of how NMR structures of MPs differ from crystal structures is the question of how to distinguish native-like from non-native MP structures. Zhou & Cross^13^ derived guidelines to evaluate native-like MP structures and suggested criteria such as hydrophobic match, small lipid exposure of hydrophilic groups, few or non-perturbing crystal contacts, oligomeric symmetry, few water or hydrophilic organics in the hydrophobic environment, and small or non-existent H/D exchange signals of amide protons in the TM region. They also suggested tight packing with few cavities as a feature of native-like structures, however, MPs require some flexibility to carry out their function. Since Cross and co-workers considered mostly crystal structures when developing these guidelines and we have shown above that crystal structures are more tightly packed than NMR structures, we argue that the tight packing requirement might originate in the differences between solution and a crystal environment.

### Quality of NMR structures can be improved via high-resolution refinement

Since we now have metrics to identify differences between NMR and crystal structures and have identified non-native features in NMR structures, we tested whether we could computationally refine them to improve some of these features. Recently, two independent groups have shown that refinement can yield more biologically relevant models when the NMR structure contains distortions that are not native-like^38^: either the refined NMR structure^39^ (MD simulations without NMR restraints) or *ab initio* Rosetta models^40^ of the single span MP KCNE1 (PDBID 2K21^41^) were docked into a potassium channel KCNQ1 model. The generated models could explain activation characteristics that have been validated experimentally.

In another study^17^, the NMR structures of soluble proteins were refined to improve stereochemistry and RMSDs compared to the crystal structure. The refined models were also better suited for phasing crystal structures. In that study, the authors used NMR restraints during refinement^42,43^, however, these proteins were in solution. Here, we abstained from the use of NMR data because: (1) We wanted to remove artifacts that stem from an artificial micelle environment, but instead want to create protein models that are consistent with a planar membrane bilayer. For DAGK for instance, the Q-value between experimental and back-calculated RDCs is much smaller for the NMR structure (Q=0.03) than for the crystal structure (Q=0.55), indicating that the large bend angle stems from the artificial micelle environment. Refining NMR structures with this artificial data would not produce models compatible with a planar bilayer. (2) Further, especially helical MPs suffer from low spectral quality (line broadening, etc.) due to the addition and influence of detergent, which makes restraint collection more challenging ^44,45^ with conceivably lower-quality restraints and higher error rates; we did not want to negatively bias refinement towards this data; and (3) we wanted to test the simplest protocol.

We refined the top NMR conformer and created 1,000 models (see Methods), the top-scoring model of which was evaluated with the metrics outlined above. Comparisons of the NMR model, the top-scoring refined NMR model, the refined NMR model closest to the crystal structure (by RMSD), and the crystal structure, are shown in Figures 5 and Supplemental Figure 9. Figure 6A compares the refined and unrefined NMR structures to the crystal structures. The refined NMR structures have smaller differences (RMSDs) to the crystal structure than the unrefined ones, illustrating reduced splaying (Figure 6B). The refined models also have improved stereochemistry as seen from lower MolProbity scores (Figure 6C) and are more tightly packed than the unrefined ones (Figure 6D). These data show that even a simple refinement protocol can nudge NMR structures of MPs toward smaller deviations from the crystal structures. These results suggest that high-resolution refinement provides a simple means to improve models of MP NMR structures and ultimately lead to a better understanding of MP structure, function, and development of disease.

**Figure 5.**
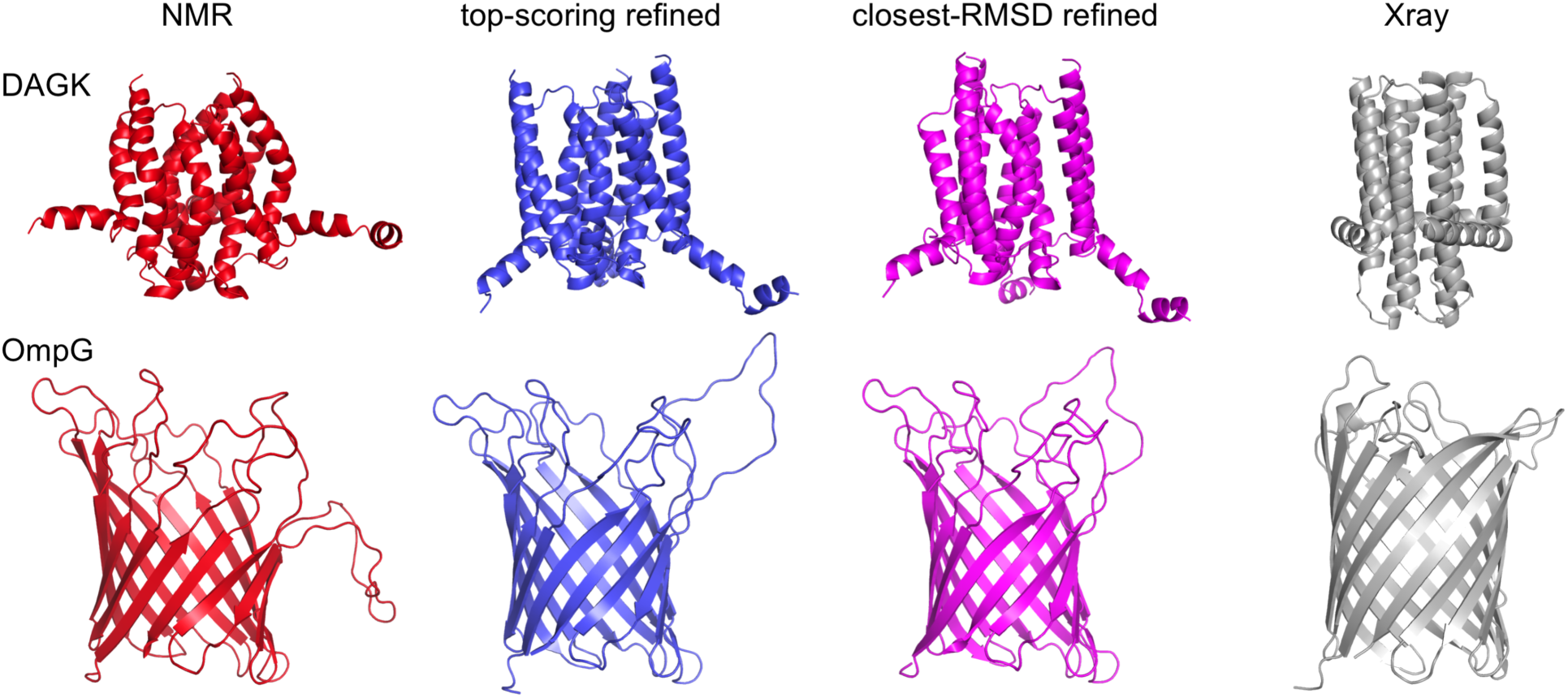
Effect of high-resolution refinement in a membrane bilayer onto structural quality. The top NMR models were subjected to Rosetta high-resolution refinement with a membrane score function to minimize discrepancies between the NMR and crystal structures. Structural features arising from NMR structure determination (see Results and Discussion) and/or membrane mimetics could be reduced: Refinement straightens out the helices in DAGK and the ‘flexible’ loop in *Yersinia pestis* OmpX that wraps around the imaginary micelle in which the structure was determined.

**Figure 6.**
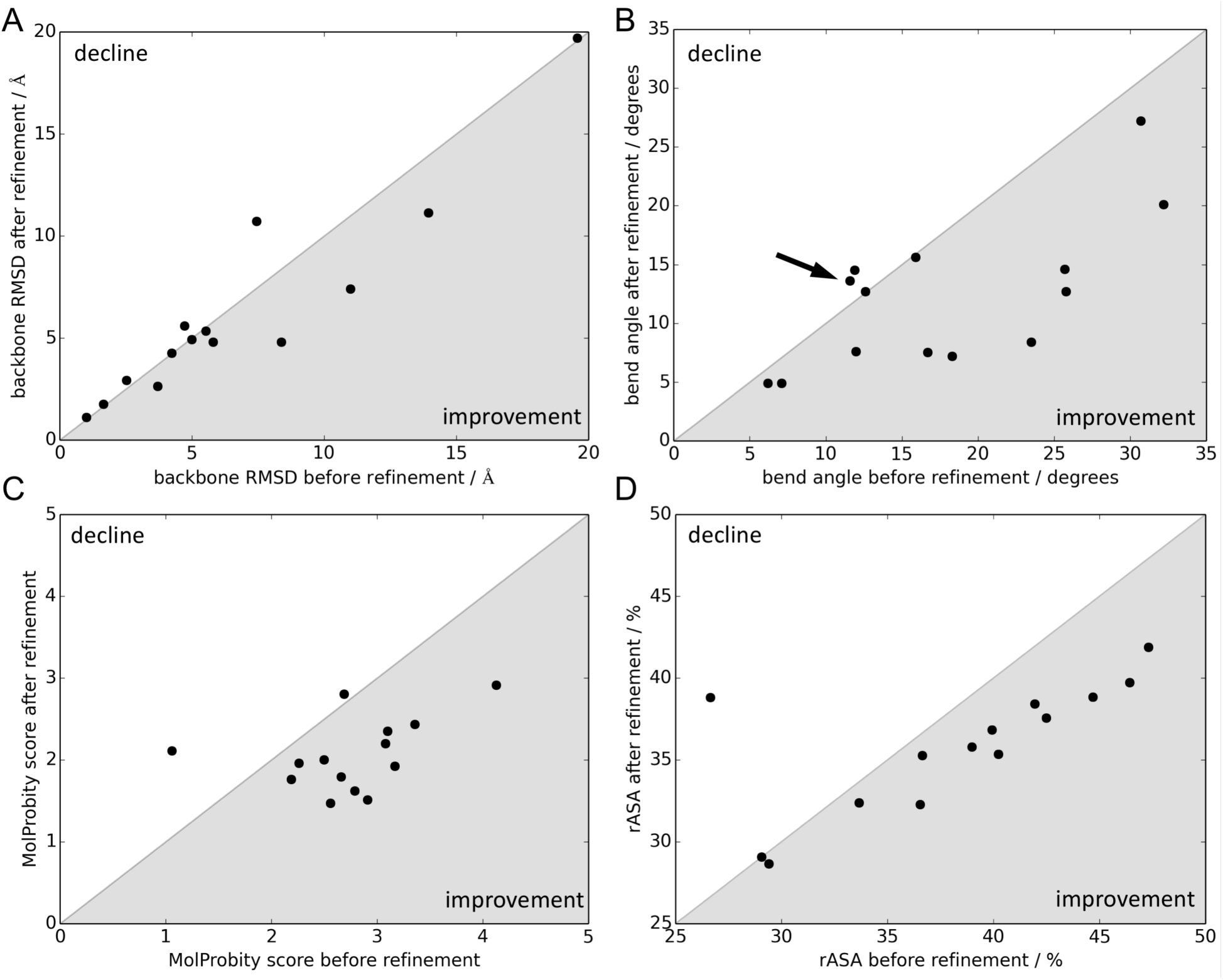
High-resolution refinement improves the quality of NMR structures. Correlation plots of structural measures before and after refinement of the NMR structure, using the top-scoring model, for the 14 membrane proteins in the dataset. Gray shaded areas denote improvements toward the crystal structure. (A) Backbone RMSD of full-length structures compared to the crystal structure. Refined NMR structures are more similar to crystal structures than unrefined ones. The outlier is bovine rhodopsin, which unfolds during the simulation due to a low-quality starting structure (Figure 1 A.6). (B) In most cases, bend angles decrease, straightening the TM region in the membrane bilayer. (C) Overall MolProbity score decreases (improves) due to an improvement in clashes, Ramachandran dihedrals and rotamers. The outlier belongs to *E. coli* OmpX, whose clash score and hence MolProbity score worsens during refinement. (D) Relative ASA averaged over all residues indicating tighter packing of refined models. The outlier is again bovine rhodopsin, which unfolds during the simulation due to a low-quality starting structure.

### Structure determination method influences structural quality

The differences between NMR and crystal structures of MPs may arise from a combination of differences in (1) structure determination methods, (2) environment of a packed crystal versus solution, (3) the hydrophobic environment of the membrane bilayer, (4) the membrane mimetic, and (5) specific lipid/detergent types used^46^. Based on the data presented here and in the literature, we believe that the strongest influence stems from different environments in a crystal vs. in solution, the method of structure determination, and membrane mimetics (see Table 2 for summary).

Given the structural differences outlined above, a natural question is: Which structure is more native-like, the crystal or the NMR structure? Since measurement of a ‘true’ structure of a MP in a native membrane bilayer remains difficult, we cannot answer that question conclusively. The structural biology community has typically used crystal structures as the standard, which might be because of the larger number of crystal structures (only about 9% of the PDB are NMR structures), the notion that proteins crystallize at an energetically favorable state, and direct detection of the electron densities (in contrast to structure calculations from NMR restraints). Even though crystal structures do not report on dynamics and can have artifacts from crystal contacts, crystal structures are typically used for method development and benchmarking: methods such as MolProbity^24^, the Dunbrack rotamer library^47^, and parts of the high-resolution Rosetta score function^48^ were derived from high-resolution crystal structures. It is therefore somewhat expected that MolProbity reports better stereochemistry for the crystal structures in our dataset and that Rosetta’s high-resolution refinement protocol creates models that are more similar to the crystal structure than to the starting NMR structure. Still, it is encouraging that a simple refinement method combined with a reliable score function can mitigate features in NMR structures that are inherent in the method, the environment, or stem from membrane mimetics. The protocol is applicable to refine NMR structures of MPs, which currently make up about 11% of the MP structures in the PDB.

Many questions remain unanswered: The impact of membrane mimetics remains somewhat elusive; the effect of different types of lipids, detergents and their mixtures on the structures are still unknown, as are the structural changes induced by other environmental variables such as pH, buffer conditions and salt concentration. Other open questions include the influence of the native environment and membrane remodeling such as curvature or lipid composition on MP structure.

## Conclusion

In this work, we have examined the differences between MP structures determined by X-ray crystallography and NMR spectroscopy, discussed the origins of these differences and tested an efficient computational refinement method to minimize them.

In addition to convergence in the NMR ensemble, which measures a combination of restraint satisfaction and the number of available restraints, we have provided four orthogonal measures of structural quality: (A) the RMSD between crystal and NMR structure describes the differences between the two, (B) the bend angle measures visible, large-scale deformations, (C) the MolProbity score defines subtler measures such as clashes, dihedral angles and rotamers that aren’t necessarily immediately visible, and (D) the overall relative accessible surface area (rASA) defines packing.

We found that TM regions of NMR ensembles typically differ within 5 Å, which is about the same range as the differences between NMR and crystal structures. These ranges are larger than what was found for soluble proteins^10-12^. Crystal structures usually have straighter TM regions and are more tightly packed than NMR structures. Crystal structures also have enhanced stereochemistry, as shown by smaller MolProbity scores. However, while crystal structures have fewer clashes than NMR models, those that exist are typically more severe. We have discussed the origins of these differences considering the different techniques by which the structures were determined, the environment of a packed crystal vs. solution, and have examined the influence of the membrane mimetics onto the structure. With this quantitative description of structural differences, we then used Rosetta high-resolution refinement on the NMR structures to try to minimize them. Our results show that refinement of the NMR structures leads to models that are more similar to the crystal structures in terms of RMSD, straightness in the TM region, stereochemistry, and packing. The protocol we present therefore provides an excellent way to improve structural quality while reducing artificial features from structure determination.

## Experimental procedures

### Dataset generation

We manually curated the PDB to find membrane proteins that had both crystal and NMR structures and found 14 proteins that match this criterion. All proteins had more NMR or crystal structures than the single pair. For crystal structures, we only added the highest resolution structures to our dataset. For NMR structures, we only considered solution state structures, since solid-state structures only exist for M2 and one for DsbB. We then manually identified the ‘best’ representatives for a pair of structures (NMR and crystal structure), given the following considerations: (1) If possible, the structures should be from the same organism (not the expression system); (2) no structural intermediates, for instance for Bacteriorhodopsin ~100 structures exist in the PDB to date, some of which are from intermediates; (3) high convergence for NMR ensembles as visually identified; (4) large number of resolved residues or similar number of residues in both structures; (5) small number of mutations between the sequences in the pair. Details about structural and sequence alignments can be found in the Supplement. The final dataset for the pairs of structures is shown in Table 1 and Figure 1.

### RMSD calculations

**RMSD between Xray and NMR:** From the processed PDBs (see Supplemental Experimental Procedures: Structural and sequence alignments) we created trimmed models which only contain the residues that are present in both structures of the pair – this means that both structures have the exact same number of residues. This allows for direct comparison between the structures and the quality measures computed for them. Structure files were then ‘cleaned’, i.e. HETATMs removed, one occurrence of seleno-methionine was mutated back to its original isoleucine, and the residues were renumbered consecutively for input into Rosetta. Rosetta ‘span files’ that contain the TM region of the protein were computed from the trimmed PDBs using the mp_span_from_pdb application^49^. Note that these structures are already embedded into the membrane bilayer as described in the Supplement. Backbone RMSDs were then computed for *all* backbone atoms in the trimmed structures, irrespective of whether the align routine had thrown them out. This was accomplished with the *rms_cur* routine in PyMOL which computes the RMSD without additional fitting or superposition. For the TM region backbone RMSDs, only the backbone atoms of the residues in the span file were used.

**RMSDs within NMR ensemble:** To compute RMSDs within the NMR ensemble, we used the models containing *all* residues; not the trimmed structures. Structure files were ‘cleaned’, i.e. HETATMs removed, one occurrence of seleno-methionine was mutated back to its original isoleucine, and renumbered consecutively for input into Rosetta. We used PyMOL’s align routine with its 5 default cycles to find an optimal superposition of the models. Backbone RMSDs were computed with the *rms_cur* routine on *all* backbone atoms in the NMR structures, irrespective of whether the align routine had previously discarded them. Rosetta span files were computed from the first NMR model in the PDB using the *mp_span_from_pdb* application^49^. For the TM region backbone RMSDs, only the backbone atoms of the residues in the span file were used.

### Computing quality measures

**TM bend angles:** Bend angles were defined as displayed in Figure 3B. The TM regions in the span file were used to define three ‘layers’ as start, center, and end residues of the TM spans (z > 0, z ~ 0, z < 0), from which the center-of-masses for each layer were computed. From these, the average radii of the TM spans were calculated, the difference of which were used to compute the bend angles. The default thickness of the hydrophobic layer was 30 Å.

**MolProbity for structural quality:** Molprobity^24^ was downloaded from https://github.com/rlabduke/MolProbity and invoked by */MolProbity/cmdline/multichart <PDB file>.* The output is written into html format which was analyzed by Python scripts. Details about how per-residue clash scores were computed in Rosetta, are given in the Supplement.

**Naccess for relative surface areas:** Relative per-residue surface areas were computed with the Naccess program^25^ with a probe size of 2.0 Å that is representative of a methyl group from a lipid acyl chain. The command line used was *naccess protein.pdb -p probe_size -r vdw.radii -s standard.data*

### High-resolution refinement

Rosetta high-resolution refinement was carried out on the trimmed structures with a recently developed MPRangeRelax algorithm (manuscript in preparation, see Supplement). This protocol carries out small dihedral angle perturbations and up to three cycles of rotamer refinement and minimization of the protein.

Details and command lines can be found in the Supplement. From 1,000 built models for each NMR structure (only MODEL 1 was considered), the top-scoring models were used for evaluation. Backbone RMSDs to the trimmed crystal structures were automatically computed during the refinement and output into the score file. TM region backbone RMSDs were computed with the rms_cur routine in PyMOL as described above.

## Author contributions

YB worked on the database, ARD analyzed the data, JKL conceived the idea, worked on the database, analyzed the data and wrote the paper with help from JJG.

## Acknowledgements

The authors would like to thank Meera Valliath for work on the database, Dr. Georg Künze for the calculation of the Q-factors for DAGK, and RosettaCommons for helpful suggestions. Funding was provided by RosettaCommons to JKL and NIH R01 Gm-078221 to JJG and JKL.

